# Unidirectional ion transport mechanism of a light-driven chloride pump revealed using X-ray free electron lasers

**DOI:** 10.1101/2021.09.21.461290

**Authors:** Toshiaki Hosaka, Takashi Nomura, Minoru Kubo, Takanori Nakane, Luo Fangjia, Shun-ichi Sekine, Takuhiro Ito, Kazutaka Murayama, Kentaro Ihara, Haruhiko Ehara, Kazuhiro Kashiwagi, Kazushige Katsura, Ryogo Akasaka, Tamao Hisano, Tomoyuki Tanaka, Rie Tanaka, Toshi Arima, Ayumi Yamashita, Michihiro Sugahara, Hisashi Naitow, Yoshinori Matsuura, Susumu Yoshizawa, Kensuke Tono, Shigeki Owada, Osamu Nureki, Tomomi Kimura-Someya, So Iwata, Eriko Nango, Mikako Shirouzu

**Affiliations:** RIKEN Center for Biosystems Dynamics Research, Yokohama, Kanagawa, Japan; RIKEN SPring-8 Center, 1-1-1 Kouto, Sayo-cho, Sayo-gun, Hyogo, 679-5148, Japan; Department of Life Science, Graduate School of Science, University of Hyogo, Hyogo 678-1297, Japan; MRC Laboratory of Molecular Biology, Cambridge Biomedical Campus, CB2 0QH, Cambridge, UK; Department of Biological Sciences Graduate School of Science, The University of Tokyo, Tokyo, 113-0034, Japan; Department of Cell Biology, Graduate School of Medicine, Kyoto University, Yoshidakonoe-cho, Sakyo-ku, Kyoto, 606-8501, Japan; Division of Biomedical Measurements and Diagnostics, Graduate School of Biomedical Engineering, Tohoku University, Sendai 980-8575, Japan; Atmosphere and Ocean Research Institute, The University of Tokyo, Tokyo, Japan; Japan Synchrotron Radiation Research Institute, 1-1-1, Kouto, Sayo-cho, Sayo-gun, Hyogo, 679-5198, Japan; Institute of Multidisciplinary Research for Advanced Materials, Tohoku University, 2-1-1 Katahira, Aoba-ku, Sendai 980-8577, Japan

**Author notes:** **To whom correspondence may be addressed. E-mail: Mikako Shirouzu**, **Eriko Nango**. **Author Contributions:** T.Ho., T.K.-S., S.I., E.N., and M.Sh. designed the research; T.Ho. and S.Y. prepared protein samples and microcrystals; T.No. and M.K. performed time-resolved visible absorption spectroscopy and analyzed the spectral data; T.No., M.K., and S.O. contributed to the pump laser setup; T.Ho., L.F., S.S., T.I., K.M., K.I., H.E., K.Kas., K. Kat., R.A., T.Hi., T.T., R.T., T.A., A.Y., M.Su., H.N., Y.M., and E.N. performed data collection; T.Na. and O.N. processed data; T.Ho., L.F., and E.N. refined the intermediate structures; K.T. developed the SFX systems at SACLA; and T.Ho., T.No., M.K., T.Na., and E.N. wrote the paper with input from all authors. Data deposition: The coordinates and structure factor for resting state, 1 ms after photoactivation state, and anion-depleted form of NM-R3 have been deposited in the Protein Data Bank, www.wwpdb.org (PDB ID code 7VGT, 7VGU, and 7VGV).

**Keywords:** microbial rhodopsin, serial femtosecond crystallography, chloride ion pump

## Abstract

Light-driven chloride-pumping rhodopsins actively transport anions, including various halide ions, across cell membranes. Recent studies using time-resolved serial femtosecond crystallography (TR-SFX) have uncovered the structural changes and ion transfer mechanisms in light-driven cation-pumping rhodopsins. However, the mechanism by which the conformational changes pump an anion to achieve unidirectional ion transport, from the extracellular side to the cytoplasmic side, in anion-pumping rhodopsins remains enigmatic. We have collected TR-SFX data of *Nonlabens marinus* rhodopsin-3 (NM-R3), derived from a marine flavobacterium, at 10 μs and 1 ms time-points after photoexcitation. Our structural analysis reveals the conformational alterations during ion transfer and after ion release. Movements of the retinal chromophore initially displace a conserved tryptophan to the cytoplasmic side of NM-R3, accompanied with a slight shift of the halide ion bound to the retinal. After ion release, the inward movements of helix C and helix G and the lateral displacements of the retinal block access to the extracellular side of NM-R3. Anomalous signal data have also been obtained from NM-R3 crystals containing iodide ions. The anomalous density maps provide insight into the halide binding site for ion transfer in NM-R3.

**Significance:** Light-driven chloride pumps have been identified in various species, including archaea and marine flavobacteria. The function of ion transportation controllable by light is utilized for optogenetics tools in neuroscience. Chloride pumps differ among species, in terms of amino acid homology and structural similarity. Our time-resolved crystallographic studies using X-ray free electron lasers reveal the molecular mechanism of halide ion transfer in a light-driven chloride pump from a marine flavobacterium. Our data indicate a common mechanism in chloride pumping rhodopsins, as compared to previous low temperature trapping studies of chloride pumps. These findings are significant not only for further improvements of optogenetic tools but also for a general understanding of the ion pumping mechanisms of microbial rhodopsins.

## Introduction

Microbial ion-pumping rhodopsins are integral membrane proteins that actively transport ions across membranes upon light stimulation (1). Bacteriorhodopsin (bR) and halorhodopsin (HR) are well-known microbial ion-pumping rhodopsins found in halophilic archaea (2, 3). bR is a light-driven outward proton pump and HR is a light-driven inward anion pump, specific for chloride ion. Microbial ion-pumping rhodopsins possess common structural features consisting of seven α-helices with an all-*trans* retinal covalently bound to a lysine residue as the chromophore, despite the transport of different ions (4). The retinal undergoes photoisomerization from the all-*trans* to 13-*cis* configuration, which initiates the photocycle accompanied by several intermediates to export ions (4, 5). Its light-controllable function is suitable for optogenetics applications for manipulating cells, such as neurons, by changing the ion concentration inside or outside the membrane (6, 7). In fact, microbial ion-pumping rhodopsins, including channelrhodopsin and HRs, are employed as optogenetic tools (8-10).

*Nonlabens marinus* rhodopsin-3 (NM-R3) is a light-driven chloride pump recently discovered in a marine flavobacterium (11). It is a distinct chloride pump from HRs and shows low amino acid sequence homology with HRs (11). To date, HR-type chloride pumps have been found in haloarchaea, marine bacteria, and cyanobacteria, including *Halobacterium salinarum, Natronomonas pharaonis*, and *Mastigocladopsins repens*, with sequence identities of 20%, 21%, and 20% to NM-R3, respectively (3, 12-15). Interestingly, NM-R3 has higher sequence identity (36%) to Krokinobacter rhodopsin 2 (KR2), a sodium pump found in *Krokinobacter eikastus* (16). NM-R3 possess a unique NTQ motif (Asn98, Thr102, Gln109) in the third helix (helix C), which corresponds to key residues (DTD motif, Asp85, Thr89, Asp96) for proton transport in bR (11, 17, 18) (Table S1). Asp85 acts as the primary proton acceptor of bR from the protonated Schiff-base (PSB), with assistance from Thr89 and Asp96, which is the proton donor (5, 17, 18). HRs from haloarchaea have a highly conserved TSA (Thr, Ser, Ala) motif, while the Ala residue is replaced by Asp in HR from cyanobacteria (19). In the X-ray crystal structure of NM-R3 (Fig. S1A), a chloride ion located between the PSB and Asn98 (Fig. S1B) is stabilized by the positive charge of the PSB (20). The position of this chloride ion is similar to those in the *Halobacterium salinarum* HR and *Natronomonas pharaonis* HR (*Np*HR) structures except for Thr and Ser, which correspond to Asn98 and Thr102 in NM-R3, respectively (20-22). Several amino acid residues around the retinal, including Arg95, Trp99, Trp201, and Asp231, are highly conserved among ion-pumping rhodopsins. Previous spectroscopic studies suggested that NM-R3 displays a similar sequence of intermediates, with K-, L-, N-, and O-like species, as in other HRs (23) (Fig. 1A). Recently, intermediate structures of NM-R3 obtained by low temperature trapping X-ray crystallography and serial femtosecond crystallography (SFX) have been reported (24, 25). However, the detailed ion-pump mechanism still remains unclear, due to the lack of dynamic structures of anion transport at atomic resolution.

**Figure 1.**
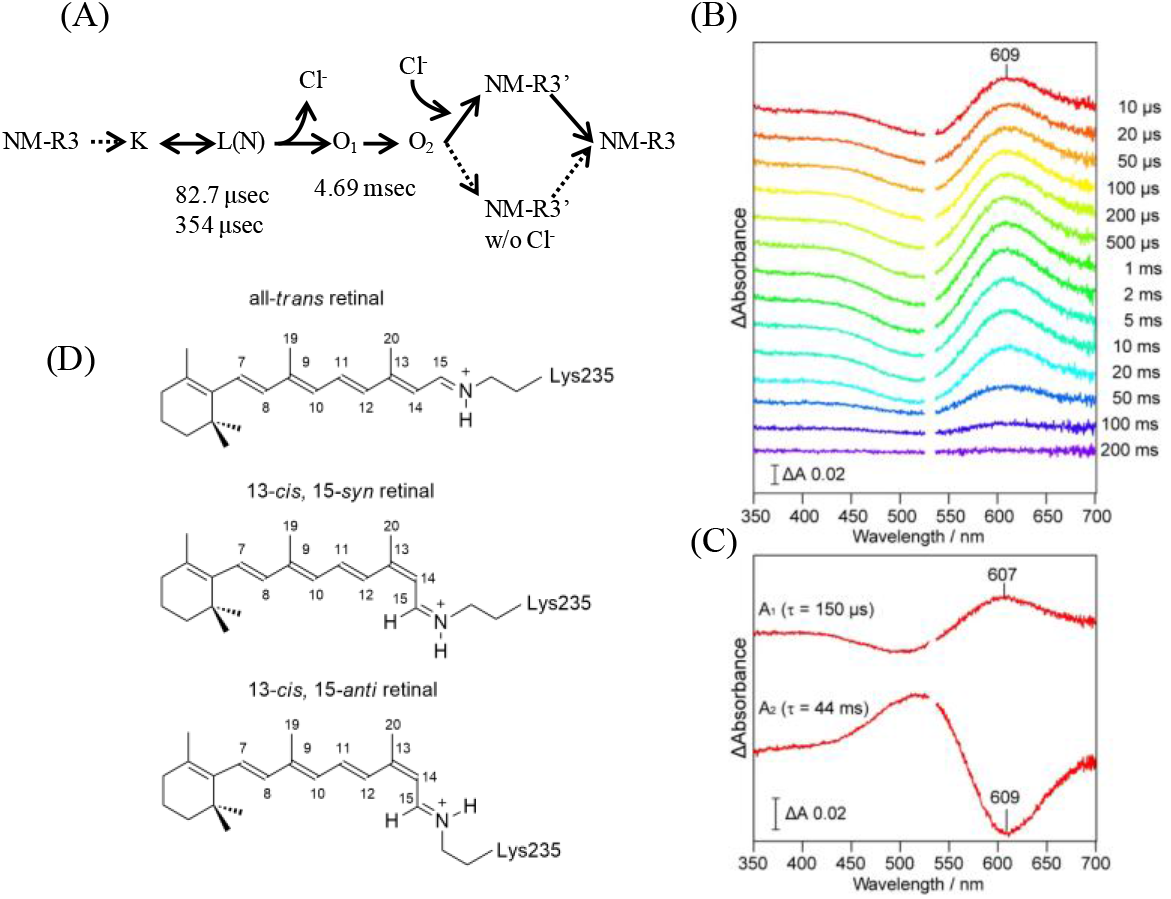
TR-visible absorption spectroscopy for microcrystals. (A) Photocycle model of NM-R3 in the 1 M NaCl buffer solution (23). (B) TR difference spectra ΔA upon the 532 nm excitation. The difference was calculated by subtracting the spectrum of NM-R3. (C) Global fitting analysis with two exponentials. The A_1_ and A_2_ amplitude spectra correspond to the differences of [ΔA_O_ – ΔA_10 μs_] and [ΔA_200 ms_ − ΔA_O_], respectively. Here, ΔA_O_ represents the difference spectrum of the O-intermediate minus NM-R3. (D) The isomeric forms of the retinal chromophore in bacterial-type rhodopsins.

Time-resolved serial femtosecond crystallography (TR-SFX) is a powerful tool for visualizing reactions and motions in proteins at the atomic level (26-28). In SFX, myriads of microcrystals are continuously injected by a sample injector into an irradiation point of X-ray free electron lasers (XFEL) at room temperature, thus providing diffraction patterns before the onset of radiation damage by the intense X-ray pulse. Combined with a visible-light pump laser for reaction initiation, TR-SFX has been applied to light-driven ion pumps to observe the structural dynamics during the ion transfer. While TR-SFX has revealed femto-to-millisecond structural dynamics in light-driven cation pumps, including bR and KR2 (29-31), TR-SFX studies of anion pumps have been limited to early-stage structures adopted at picoseconds after light illumination (32). In addition, although NM-R3 pumps a chloride ion (Cl^-^) as a physiological substrate, it can also transport bromide (Br^-^), iodide (I^-^) and other anions from the extracellular side to the cytoplasmic side (23). I^-^ or Br^-^ serves as a marker for tracking the positions of ions, due to the greater number of electrons, whereas Cl^-^ is less distinguishable in X-ray crystallography. Therefore, TR-SFX experiments using I^-^ or Br^-^ are expected to directly visualize the process of ion transport.

Here, we report the conformational alterations in NM-R3 during Br^-^ or I^-^ pumping, obtained by both TR-SFX and time-resolved spectroscopy of crystals. The resulting sequence of movements in NM-R3 demonstrates how the chloride pump transports anions with a large ionic radius and prevents the backflow of anions from the cytoplasmic side.

## Results

### XFEL Structures of bromide-bound or iodide-bound NM-R3 in the resting state

In 2016, we reported the crystal structure of NM-R3, determined at 1.58 Å resolution from data collected using synchrotron radiation (SR) (20). The X-ray structure of NM-R3 from crystals grown in a lipidic cubic phase (LCP) indicated that one Cl^-^ is bound to the PSB of the all-*trans* retinal (Fig. S1). That same year, Kim et al. also determined the X-ray structure of NM-R3 at 1.56 Å resolution using SR, which showed two Cl^-^ : one at the same position and the other by the loop between helix A and helix B on the cytoplasmic side (33). To date, several structures of NM-R3, including the structures from NM-R3 crystals obtained at lower pH and the bromide-bound NM-R3, have been reported (33). Recently, the XFEL structure of chloride-bound NM-R3 was also solved at the Linac Coherent Light Source (25).

We have determined the structures of the bromide-bound and iodide-bound forms of NM-R3 crystals grown in LCP at 2.1 Å resolution, using XFEL pulses at the SPring-8 Angstrom Compact Free Electron Laser (SACLA) (Fig. S2A). The dissociation constants (*K*_d_) for Cl^-^, Br^-^ and I^-^ in NM-R3 were previously estimated as 24 mM, 10 mM and 2.5 mM, respectively, by Tsukamoto et al. (23). At first, NM-R3 crystals embedded in LCP were soaked in a solution of 1 M sodium iodide to prepare the iodide-bound form, and then injected from a high-viscosity sample injector into the XFEL irradiation area (34). However, the interaction of the sample with XFEL emitted light and the sample stream flow was very unstable and formed droplets, presumably due to the phase transition of LCP. Since the highly concentrated halide ions caused liquefaction of sample streams, the concentration used for soaking needed to be decreased to the extent that the phase transition and illumination of the samples subsided. Consequently, NM-R3 crystals soaked in buffer containing 200 mM sodium iodide or 1 M sodium bromide were successfully injected, without phase transition and illumination.

The overall XFEL structures of the bromide-bound NM-R3 (NM-R3-Br) and iodide-bound NM-R3 (NM-R3-I) are similar to the SR structure of the chloride-bound NM-R3 (NM-R3-Cl) (PDB ID: 5B2N), with Cα rmsds of 0.41(Br) and 0.32(I) Å (20) (Fig. S3). Although the number of water molecules was decreased from 96 to 53, most of the internal water molecules were consistent with those in the SR structure. Three halide ions were detected in both the NM-R3-Br and NM-R3-I XFEL structures (25) (Fig. S2). The first halide ion is bound to the PSB, as in the NM-R3-Cl SR structures (Fig. S4A-C). The second halide ion is in the loop between helix A and helix B in both the NM-R3-Br and -I structures, as in the NM-R3-Cl SR structure reported by Kim et al. (Fig. S2B) (33). The third halide ion is on the cytoplasmic side of helix E (Fig. S2C).

In contrast, the distances between the anion and the key residues around the retinal differ in the NM-R3-Br and -I XFEL structures, as compared with the SR structures (NM-R3-Cl and NM-R3-Br). Br^-^ and I^-^ are bound to the PSB with 3.18 and 3.37 Å distances, respectively, depending on their ionic radii (Fig. S4A and B). Interestingly, the distances between Br^-^ (or I^-^) and two residues (Thr102, Asn98) are relatively shorter or almost the same as those of the SR structures (Fig. S4C and D). Particularly, the interaction with Asn98 is tightened, with distances of 3.10 (Br-) and 3.30 (I-) Å, as compared to that of NM-R3-Cl (4.04 Å) (Fig. S4A-C). The NM-R3-Cl XFEL structure (5ZTL) reported previously also indicates this tendency (Fig. S4E) (25). As a significant difference, only one water molecule is bound to the halide ion in our XFEL structures, as in the other NM-R3 SR structures, whereas an additional water molecule bound to Asp231 was observed in 5ZTL (25).

### Time-resolved spectroscopy analysis of NM-R3 microcrystals

Previously, Tsukamoto et al. reported that the photocycle of NM-R3-Cl in a solution including DDM consists of five kinetically defined states: K, L(N), O_1_, O_2_, and NM-R3’, as photointermediates determined by TR visible absorption spectroscopy (Fig. 1A) (23). The two O-intermediates are specific to NM-R3, since a single O-intermediate is seen in other HRs (18). In the photocycle model, Cl^-^ is released during the transition from L(N) to O_1_ (23). Subsequently, a conformational change may occur to prevent the reverse ion transfer and to prime the next uptake of Cl^-^ from the extracellular side during the O_1_-O_2_ transition.

In this study, we investigated the photocycle kinetics of NM-R3-Br in the crystalline phase by TR-visible absorption spectroscopy. The TR difference spectra after 532 nm excitation are shown in Fig. 1B. At 10 μs, a broad positive difference peak arises at a relatively long wavelength around 610 nm, suggesting the possibility of the partial existence of an O-intermediate together with earlier intermediates (K or L/N). The positive peak then increases with a slight blue shift with a time constant (τ) of 150 μs, and vanishes with a τ of 44 ms, as confirmed by a global fitting analysis with two exponentials (Fig. 1C). This spectroscopic result demonstrates the accumulation of the O-intermediate in NM-R3-Br microcrystals, followed by its decay to the initial state of NM-R3. It should be noted that our spectral analysis detects only one O-intermediate, while two O-intermediates, O_1_ and O_2_, have been proposed (23). Although it is difficult to identify which type of O-intermediate could be accumulated in the microcrystals, we tentatively assigned the observed species to O_1_, according to its characteristic long wavelength (>600 nm) of the positive difference peak. This assignment will be discussed further in the next section and the discussion section.

### Structural changes in NM-R3-Br crystals after photoexcitation at Δ*t*=1 ms

TR-SFX data were collected from NM-R3-Br and NM-R3-I microcrystals, using a TR-SFX setup at SACLA (35). The microcrystal stream was continuously ejected from the nozzle of the high-viscosity sample injector, and then irradiated with an XFEL pulse at 1 ms after photoactivation by a 540-nm, nanosecond laser pulse.

TR-SFX data of NM-R3 crystals with a bromide ion were recorded to 2.1 Å resolution. In the |*F*_obs_|^light^ - |*F*_obs_|^dark^ difference electron density map, most of the positive and negative features are visible around the retinal and on helices C, F, and G (Fig. S5). The paired negative and positive electron density peaks appearing around the C12-C14 and C13 methyl groups of the retinal reveal that the retinal moves toward helix C (Fig. 2A-C). The negative and positive difference features indicate that the side chain and backbone of Lys235, which is covalently attached to the retinal, are slightly displaced toward the cytoplasmic side. The time-resolved difference absorption spectra from NM-R3 crystals demonstrate that the O-intermediate (O_1_ or O_2_) is dominant at Δ*t* =1 ms (a delay time of 1 ms after photoexcitation). The re-isomerization of the retinal from the all-*trans* to 13-*cis* configuration was previously proposed to occur in the L(N) to O transition (Figs. 1D and S6) (23). In general, the O-intermediates of bacterial-type rhodopsins have been considered to adopt the all-*trans* (13-*trans*/15-*anti*) retinal configuration (5, 36, 37). However, we modeled the features around the retinal in the 13-*cis*/15-*syn* configuration, since it was more compatible with the difference electron density than the all-*trans* or 13-*cis*/15-*anti* configurations (Figs. 2B). Previous low-temperature trapping crystallographic studies of *Np*HR showed that the N-like intermediate termed X(N’), before the O-intermediate, adopts the 13-*cis*/15-*syn* configuration (37). Given that the 13-*cis*/15-*syn* configuration is in the transition from 13-*cis* to all-*trans*, this intermediate at Δ*t* =1 ms would be the O_1_-intermediate in an early stage of the O-state.

**Figure 2.**
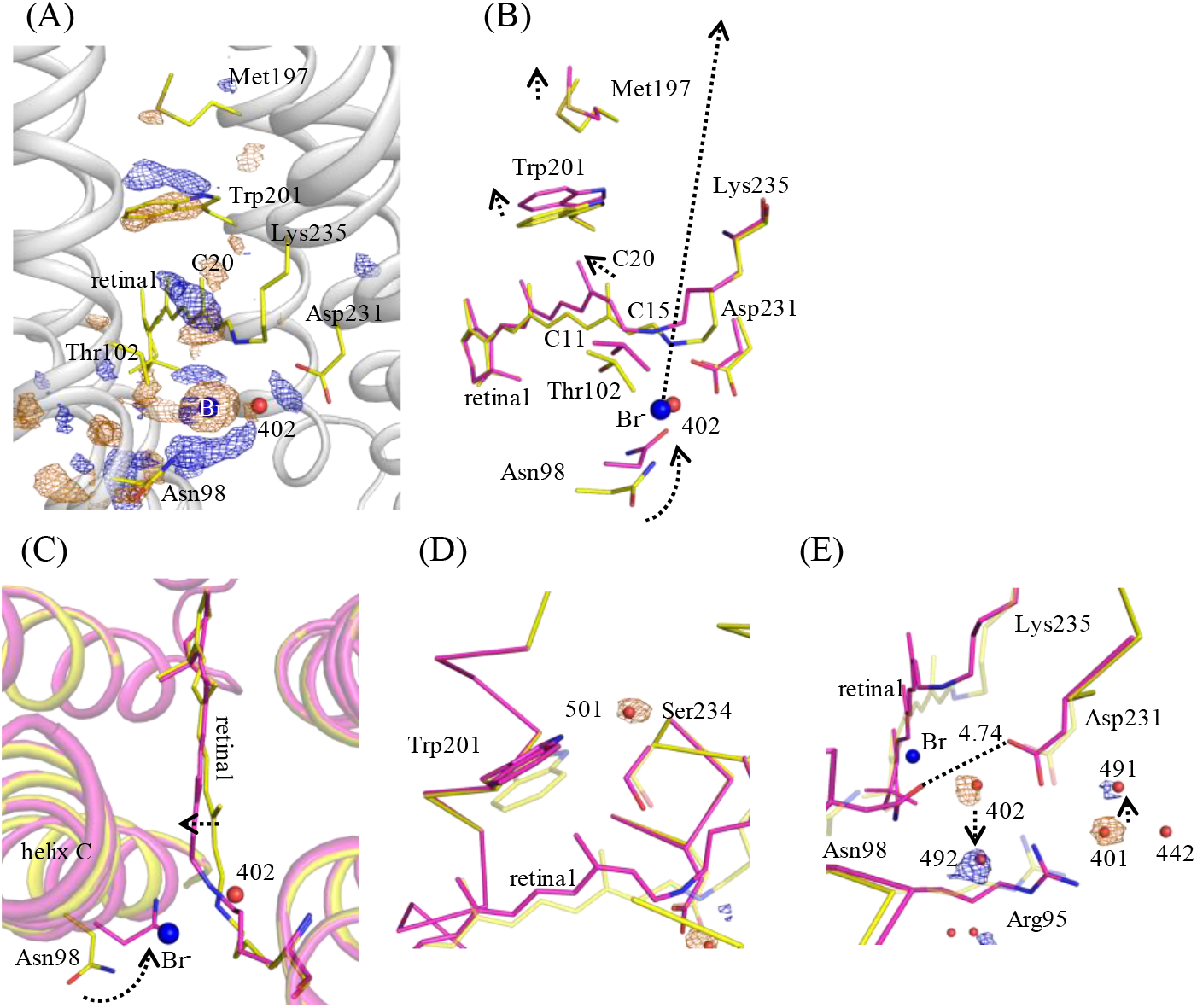
Structural change of NM-R3-Br near retinal at Δ*t* =1 ms. (A) View of the |F_obs_|^light^ - |F_obs_|^dark^ difference Fourier electron density map near the retinal in NM-R3-Br for Δ*t* =1 ms. The blue electron density map indicates positive electron density, and orange denotes negative electron density (contoured at ±3.5σ). The resting state NM-R3 model (yellow sticks and white helices) was used for phases when calculating this map. (B) and (C) Crystallographic structural models derived from partial occupancy refinement are superimposed upon the resting state NM-R3 structure (yellow) for Δ*t* = 1 ms (pink). View of the |F_obs_|^light^ - |F_obs_|^dark^ difference Fourier electron density map at the water molecules Wat501 (D) and Wat401 and Wat402 (E) for Δ*t* =1 ms. The blue electron density map indicates positive electron density, and orange denotes negative electron density (contoured at ±3.0σ). Crystallographic model for the time point Δ*t* =1 ms (magenta) superimposed on the resting state model (yellow, partially transparent). A bromide ion and water molecules are depicted by blue and red spheres, respectively. The movements of the retinal and water molecules are depicted by a dashed arrow. Numbers indicate the distance (Å) between two atoms.

On the intracellular side of the retinal, paired negative and positive difference electron density features reveal that the C13 methyl group of the retinal moves toward the cytoplasmic side by 1.70 Å, with a corresponding 1.05 Å displacement of the Trp201 indole ring in helix F (Fig. 2B). The side chain of Trp201 interacts with Wat501 in the resting state, but the displacement of Wat501 is observed at Δ*t* = 1 ms as a negative difference electron density peak (Fig. 2D). Similar movements of Trp182 in bR and Trp215 in KR2, which are genetically and structurally conserved in most bacterial-type rhodopsins, were observed in previous TR-SFX studies (29, 31). The negative and weakly positive difference electron density peaks along Met197 show that the Met197 side chain slightly moves toward the cytoplasmic side upon the sequential displacement of Trp201 and the retinal (Fig. 2A and B).

The structural model for Δ*t* = 1 ms indicates that the PSB is oriented toward the cytoplasmic side and interacts with Thr102. As a result, the distance between Thr102 and the PSB becomes closer, from 4.21 Å to 3.45 Å. The strongly negative difference electron density features around the bromide ion bound to the PSB position also reveal that the halide ion is displaced. No positive difference electron density peak comparable in size to a bromide ion is observed. The paired negative and positive electron densities associated with Asn98 of helix C indicate that Asn98 moves toward the empty space created when the bromide ion bound to the PSB is dislodged (Fig. 2B). Consequently, the Cα and side chain OD1 of Asn98 move toward the retinal by 1.76 Å and 5.23 Å, respectively (Fig. 2B). The paired negative and positive difference electron densities arising along the extracellular side portions of helix C (residues Gly93-Leu108) were modeled as an inward movement of helix C (residues Tyr94-Thr102) toward helix G (Fig. 3A and 3C). This inward flex of helix C is similar to the structural changes in the TR-SFX data of bR (29). On the other hand, the weaker paired negative and positive electron density features along the cytoplasmic side of helix C show that residues Ile103-Leu108 move slightly outward, away from the retinal (Fig. 3B). This outward movement of the cytoplasmic side of helix C could create an ion translocation pathway in the cytoplasmic interhelical space.

**Figure 3.**
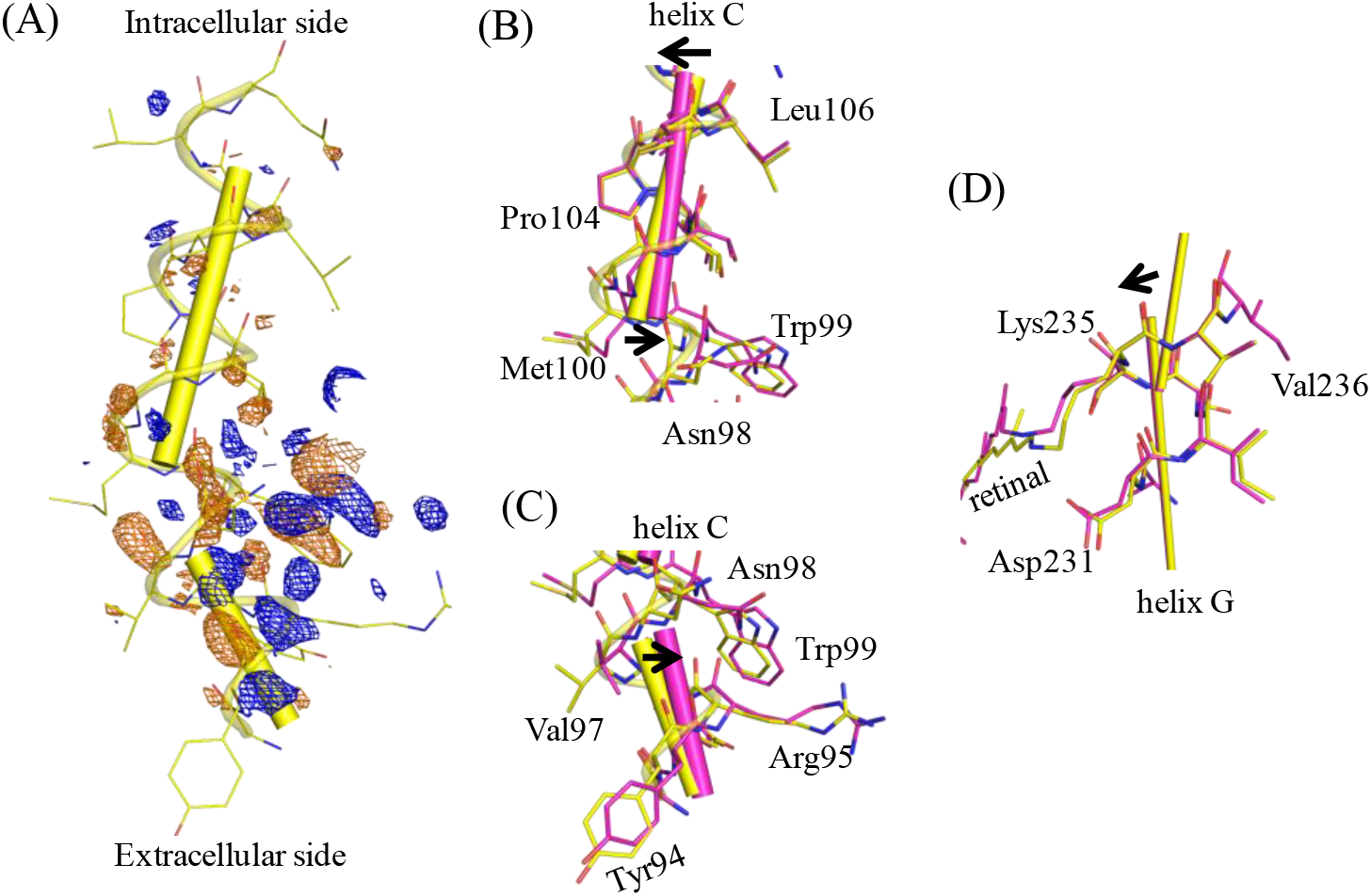
Conformational changes of helices C and G. (A) View of the |F_obs_|^light^ - |F_obs_|^dark^ difference Fourier electron density map along helix C and the resting state structural model (yellow). The blue electron density map indicates positive electron density, and orange denotes negative electron density (contoured at ±3.0σ) for Δ*t* =1 ms. (B) Structural model of helix C on the intracellular side (Asn98-Leu106) for the time point Δ*t* =1 ms (pink) superimposed upon the resting state model (yellow). (C) Structural model of helix C on the extracellular side (Tyr94-Asn98) for the time point Δ*t* =1 ms (pink) superimposed upon the resting state model (yellow). (D) Structural model of helix G near the retinal for the time point Δ*t* =1 ms (pink) superimposed upon the resting state model (yellow). The helices C and G move towards the inside of the protein in association with the movements of Asn98 and Lys235, respectively.

The paired negative and positive difference electron densities along helix G demonstrate that this helix is slightly shifted toward helix C (Fig. 3D). A positive difference electron density feature appears between Asn98 of helix C and Asp231 of helix G. We modeled the feature as a movement of the side chain of Asp231 toward Asn98 (Fig. 2B and E). The distance between Asn98 and Asp231 becomes closer (from 10.27 Å to 4.74 Å, Asn98 OD1-Asp231 OD1) with the movements of helices C and G, which block the pathway inside the protein, like a gate closing. The gate composed of the two hydrophilic amino acids may play a role in preventing back-flow or overflow of anions by forming a steric or electrostatic barrier.

A negative difference electron density peak visible on a water molecule, Wat402 between Asn98 and Asp231, indicates a displacement of Wat402 (Fig. 2E). In the resting state, Wat402 forms hydrogen-bonding interactions with the bromide ion, Asp231, Trp99, and Arg95. Correlated displacements are observed on Trp99 and Arg95. The paired negative and positive difference electron density features reveal that Trp99 moves toward the retinal. The negative difference electron density and 2 *F*_obs_ – *F*_calc_ electron density features along Arg95 show that the side chain of Arg95 is disordered by the loss of hydrogen-bonding interactions. The similar disordering of Arg95 was seen in low temperature studies of NM-R3 in SR and TR-SFX experiments with KR2 (corresponding to Arg109 in KR2) at 1 ms following photoactivation (24, 31). The equivalent arginine residue is presumedly involved in the ion transfer in bR and *Np*HR (38, 39). The disordering of Arg95 may facilitate anion uptake from the extracellular side. The sequential movements on the extracellular side trigger displacements of water molecules, detected as positive or negative difference electron density peaks, indicating the disordering of Wat401 in addition to Wat402 and the ordering of new water molecules, Wat491 and W492 (Fig. 2E). Low temperature trapping crystallographic studies of NM-R3 have shown that negative difference electron density peaks appear on Wat401, Wat402, and Wat442, but no positive electron density peak arises around the water molecules except for Wat402 (24), which is inconsistent with our observations.

### Implications for structural changes and ion transport pathway in NM-R3-I after photoexcitation at Δ*t*=10 μs and 1 ms

Bromine and iodine display anomalous dispersions in specific X-ray wavelength ranges. The K absorption edge of bromine (at 13.5 keV) was beyond the easily accessible wavelength at the XFEL facility. Moreover, such shorter wavelengths are prone to yield low resolution data due to the lower quantum efficiency of the detector, including the MPCCD detector used at SACLA (38). On the other hand, iodine has a sufficiently anomalous *f*’’ value of 8.6 e^-^ at 7 keV, which is often used at SACLA (*f*’’ value is taken from http://skuld.bmsc.washington.edu/scatter/). We attempted to measure an anomalous signal from NM-R3-I crystals to identify an anion transfer pathway, and obtained TR-SFX data at Δ*t* =10 μs and 1 ms after photoactivation. In both data sets, negative and positive difference electron density features around the retinal are visible, but become weaker as compared with the difference electron density map of the TR-SFX data of NM-R3-Br recorded at Δ*t* =1 ms. Furthermore, in the difference electron density map of NM-R3-I at Δ*t* =1 ms, very weak negative electron density peaks are observed on Wat401 (3.3σ) and Wat402 (3.3σ) and positive electron peaks corresponding to Wat491 or Wat492 are not detected, which implies the lower accumulation of a photo-intermediate in NM-R3-I. The binding affinity of I^-^ (2.5 mM) estimated from previous spectroscopic studies is higher than that of Cl^-^ (24 mM) (23). Therefore, it may cause lower ion pumping efficiency in NM-R3.

As remarkable difference electron density features, the paired negative and positive electron densities along the retinal show that the retinal moves toward helix C at Δ*t* =10 μs (Fig. S7A). Strong paired negative and positive electron density peaks appearing on the iodide ion interacted with the PSB, indicating a displacement of the iodide ion from the retinal toward Thr102 on the cytoplasmic side at Δ*t* =10 μs (Fig. S7A). The paired negative and positive electron density features around Asn98 suggest that the side chain of Asn98 in helix C also moves toward the iodide binding site (Fig. S7B), which results in the distortion of helix C (Arg95-Leu108). As with the difference electron density changes observed in the difference map of NM-R3-Br at Δ*t* =1 ms, the paired negative and positive electron density features along Trp201 show that Trp201 of helix F also shifts toward the cytoplasmic side (Fig. S7). No difference electron density peak associated with Asp231 in helix G is observed at Δ*t* =10 μs, whereas Asp231 is displaced toward the retinal in NM-R3-Br at Δ*t* =1 ms. On the other hand, the difference electron density features seen in NM-R3-I at Δ*t* =1 ms are similar to those of NM-R3-Br recorded at Δ*t* =1 ms. However, fewer difference electron density features along helices F and G are observed in NM-R3-I at Δ*t* =1 ms, while the difference electron density features associated with helices F and G clearly appear in NM-R3-Br at Δ*t* =1 ms.

Strong anomalous electron signals are visible at the three iodide ion sites observed in the resting state in both anomalous density maps for Δ*t* =10 μs and 1 ms (Fig. S8), indicating that most of the iodide ions remain with the protein. In particular, the anomalous density maps show the signals of 28.6σ for Δ*t* =10 μs and 32.5σ for Δ*t* =1 ms at the iodide binding site around the retinal. Several anomalous peaks are found on the surface of NM-R3-I, due to the high iodide ion concentration (200 mM). Very few remarkable anomalous signals (3.5σ) are visible in the solvent channel inside NM-R3-I, while anomalous densities from the sulfur atoms of Cys and Met residues inside the protein are observed (Fig. S8). At Δ*t* =10 μs, an anomalous peak (4.4σ) arises near Leu111-Leu114 of helix C (Fig. S8A) and at Δ*t* =1 ms, a small anomalous peak (4.1σ) is situated between Ile190-Val193 of helix F and Ile245 of helix G (Fig. S8B). These peaks are located at the anion release exit on the intracellular side. However, no difference electron density peaks are detected around these locations, and thus they are not a major position for I^-^ at the time points but may instead indicate part of an anion transport pathway. Low temperature trapping studies of NM-R3 have shown that an electron density peak of bromide ion is observed around Cys105, Leu107, and Leu108 in helix C (24). In addition, low temperature trapping studies of *Np*HR have suggested that a bromide ion is displaced to a temporal cavity between the PSB and the side chain of Ile134 (corresponding to Leu106 of helix C in NM-R3) (37). Given these observations, light-driven chloride pump rhodopsins may transport Cl^-^ along helix C into the cell. Other anomalous peaks are also observed within the anion uptake region on the extracellular side at Δ*t* =10 μs and 1 ms. Anomalous density peaks appear along Tyr208 of helix F at both time points (3.4σ for Δ*t* =10 μs and 4.1σ for Δ*t* =1 ms) (Fig. S8). In this case, a difference electron density peak is not visible in either of the difference maps of NM-R3-I, but these locations may be candidates for a transport pathway on the anion uptake side.

### Anion-depleted form

We also obtained a different form of NM-R3 crystals belonging to the space group *P2*_*1*_*2*_*1*_*2*_*1*_, using crystallization conditions containing Cl^-^, whereas the NM-R3 crystals used for TR-SFX are the *C2* form with one monomer per asymmetric unit. The *P2*_*1*_*2*_*1*_*2*_*1*_ crystals comprise three protomers per asymmetric unit, and their structures (chains A-C) are different from one another (Fig. S9A). Chains A and B adopt an apo form, due to the lack of Cl^-^ bound to the retinal (Fig. S9B). The side chain of Asn98 in helix C is located at the anion binding site instead of Cl^-^, in association with the displacement of Thr102 (Fig. S9B). This induces the movement of helix C toward helix G, as in the O_1_ intermediate structure obtained by TR-SFX. However, the conformations of the other helices are almost the same as those in the resting state structure, except for helix C. A previous structural analysis of the anion-depleted form of *Np*HR indicated that the side chain of Thr126 (equivalent to Asn98) moves toward the halide binding side, accompanied by the inward movement of helix C (39). Asn98 is likely to serve as a stabilizer in the absence of a halide ion. On the other hand, the retinal of chain C binds Cl^-^ and is very similar to the structure in the *C2* crystal. An overlay of the chain C structure of the *P2*_*1*_*2*_*1*_*2*_*1*_ crystal with the *C2* crystal structure reveals a rmsd of 0.49 Å for the backbone Cα atoms.

## Discussion

Low temperature trapping studies of *Np*HR have provided multiple intermediate structures, including the L_1_, L_2_, N, and O states (37, 40). The L_1_ structure indicated that Cl^-^ remains bound to the PSB, as in the resting state structure (39). In the L_2_ structure, the anion slightly moved to a position between Ser130 and the PSB, but no other movements were observed (37). Similarly, our NM-R3-I structure at Δ*t* =10 μs indicates that I^-^ moves toward Thr102, a conserved residue in the NTQ motif that is equivalent to Ser130 of the TSA motif in *Np*HR. In contrast, the Asn98 side chain moves toward the anion binding site in the resting state, in association with the large movement of helix C. Previous spectroscopic studies of NM-R3 indicated the presence of the intermediate states K, L(N), O_1_, O_2_, and NM-R3’ (23) (Fig. 1A). The NM-R3-I structure at Δ*t* =10 μs is presumed to be the L(N) state, since the K state appears at an earlier step after light illumination. Interestingly, TR-SFX studies of NM-R3 that measured early-stage dynamics (1-100 ps) showed that Cl^-^ is already translocated to Thr102 upon retinal isomerization at Δ*t* =100 ps and the C_α_ of Asn98 slightly moved within 1 Å (32), in contrast to the movements in the NM-R3-I structure at Δ*t* =10 μs. Meanwhile, Trp99 and Trp201, located around the retinal, were displaced from 1 ps to 100 ps as observed in the NM-R3-I data at Δ*t*=10 μs (32). Therefore, photoisomerization of the retinal may trigger the displacements of Cl^-^ toward Thr102 and two tryptophan residues, followed by the movement of Asn98 in helix C to fill in the space left after the Cl^-^ moved.

We assigned the NM-R3 at Δ*t* =1 ms to the O_1_ intermediate, based on the 13-*cis*/15-*syn* configuration of the retinal and the elimination of the halide ion bound to the PSB (Figs. 2B and S6). This halide ion is released to the extracellular side at Δ*t* =1 ms, since no noticeable peak is visible in the difference electron density map and the anomalous density map. Interestingly, the *Np*HR N-like (X(N’)) structure determined from low temperature trapping studies also indicated the 13-*cis*/15-*syn* configuration of the retinal and the elimination of the halide ion (37). In the *Np*HR X(N’) structure, large movements of helix C originating by the displacement of Thr126 were observed, resembling the movement of helix C including Ans98 (equivalent to Thr126) in NM-R3 at Δ*t* =1 ms. Likewise, the side chain of Thr126 in *Np*HR moved toward the space created by the elimination of the halide ion. Although NM-R3 shares low sequence identity to *Np*HR with the different key motif, they have obvious similarities in terms of the structural changes that take place during halide ion transfer. In addition, light/dark adaptation does not occur in both NM-R3 and *Np*HR, unlike the light/dark adaptation cycle of bR, which is attributed to retinal isomerization from the all-*trans* to 13-*cis* configuration (23, 41). These two chloride pumps might have a common mechanism for halide ion transfer, although NM-R3 evolved independently from *Np*HR. On the other hand, helices C and F underwent large conformational changes in the *Np*HR X(N’) structure, thus altering the ion accessibility from the cytoplasmic side to the extracellular side (22). However, in the NM-R3 O_1_ structure, the movements of helix F are smaller than those of helix C. The restricted movements of helix F might be caused by the tight crystal packing of NM-R3 in the LCP, since *Np*HR was crystallized by vapor diffusion.

Our study revealed that the Asp231 side chain moves toward the halide ion site around the PSB in NM-R3 at Δ*t* =1 ms, accompanied by the displacement of Asn98 (Figs. 2E and 3D). These movements have not been observed in previous NM-R3 studies. To our knowledge, this is the first observation of the movement of Asp231 in chloride pump rhodopsins. Previous studies of *Np*HR mutants demonstrated that the mutation of Asp252 (equivalent to Asp231 in NM-R3) has no detectable light-driven chloride ion transportation activity (42). Therefore, Asp231 may be one of important residues for anion transportation. The movement of Asp231 toward the halide binding site dislodges Wat402, thus disordering Arg95 and Wat401 (Fig. 2E). Changes in the hydrogen bonding networks facilitate anion transportation. Furthermore, the side chain carboxylate group of Asp231, with its negative charge, might serve as a key driving force for the halide ion transportation around the PSB. Asn98 and Asp231 approach each other within 5 Å by the movements of helices C and F, which block water molecules and ions from accessing the extracellular side, like closing a gate, and prevent the backflow of halide ions (Fig. 4). The gate closure by Asn98 and Asp231 is a distinctive change, supporting the alternative access model proposed by Jardetzky (43). The principle of the model is that the protein is initially accessible on one side of the membrane and switches conformations to a low-affinity binding site that is accessible to the other side. In addition, the structural model of the NM-R3-Br reveals the outward movements of helix C at Δ*t* =1 ms, which allow access to the cytoplasmic side. Interestingly, the displacement of the retinal toward helix C is observed at Δ*t* =1 ms, and thus also invokes a disconnection between the cytoplasmic and extracellular sides (Fig. 2C). This retinal movement resembles that seen in the TR-SFX study of channelrhodopsin (ChR) at Δ*t* =1 ms, by Oda et al. (44). However, helix C of ChR was pushed outward by the retinal conformational change, while helix C of NM-R3 slightly moved within 0.5 Å. The authors proposed that the sequential movements of ChR induced water influx from the cytoplasmic side (44). The structural changes of the retinal and helix C in NM-R3 may assist the anion pump and/or disrupt the pathway for the reverse anion transfer.

**Figure 4.**
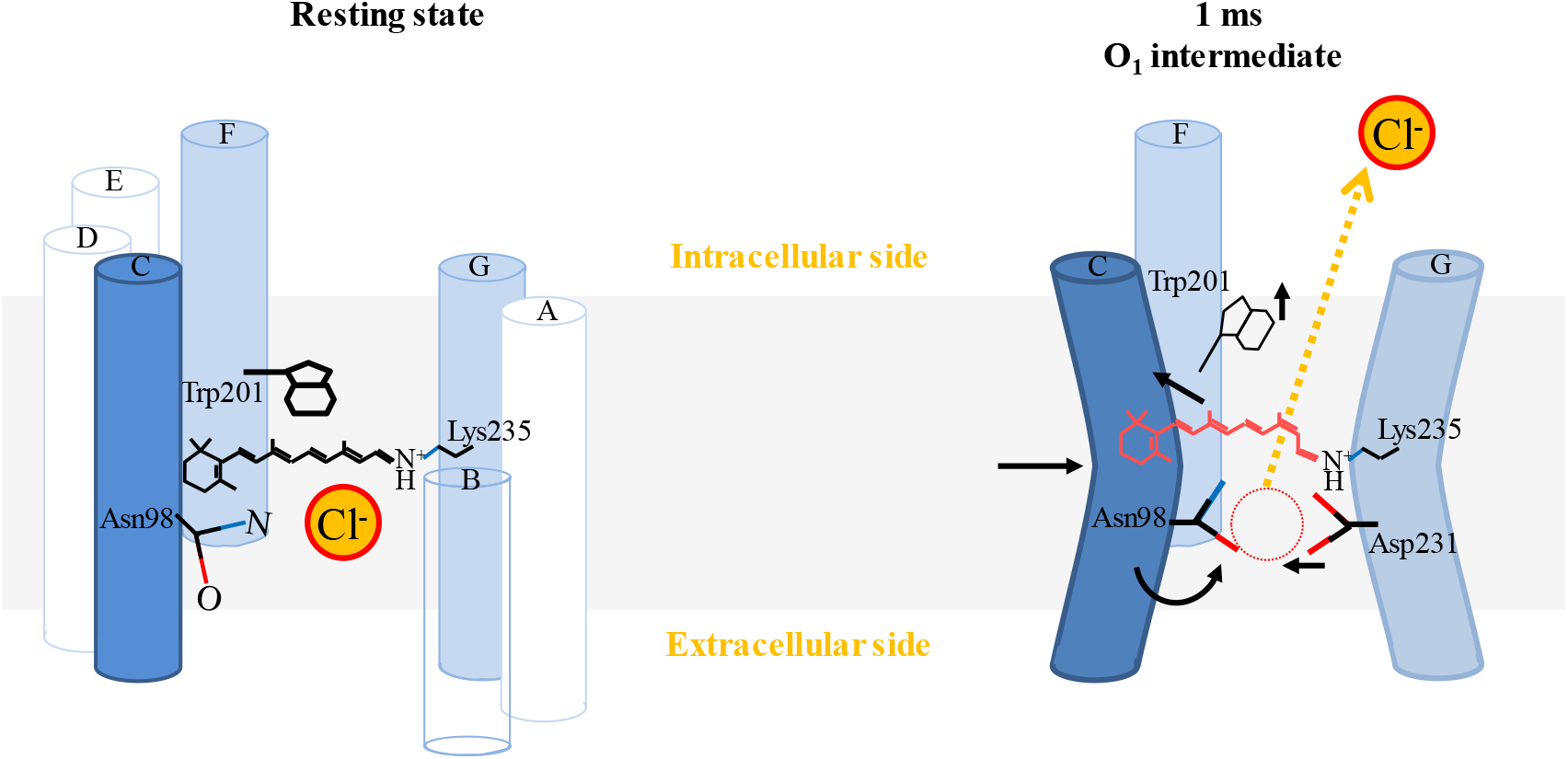
Schematic of the conformational changes during the ion transport. NM-R3 resting state structure highlighting Asn98, the anion bound to the retinal, and Trp201(left panel). Photo-isomerization of the protonated retinal from all-*trans* (black) to 13-*cis* triggers the transfer of the chloride ion (anion). At Δt =1 ms (right panel), the middle of helix C bends at Asn98 toward the inside, and Trp201 moves toward the cytoplasmic side. The retinal adopts the 13-*cis*/15-*syn* configuration (orange).

NM-R3 has high amino-acid sequence and structural feature similarities with KR2, the light-driven Na^+^-pumping rhodopsin. In particular, the key amino acid residues involved in ion transfer around the retinal are identical, except for Thr102 (Asp116 in KR2), whereas they pump oppositely charged ions. However, the two rhodopsins are different in terms of their photocycles and the initial locations of the ions in the resting state. KR2 forms the intermediate states K, K_L_, L/M, O_1_, and O_2_ during the photocycle, while the M state does not appear in the NM-R3 photocycle (31). A sodium ion is not bound within the retinal binding pocket in the KR2 resting state structures (45, 46). TR-SFX studies of KR2 showed that the sodium ion is located around the deprotonated SB, at a distance of 2.5 Å from Asn112 (equivalent to Asn98 in NM-R3) and Asp251 (equivalent to Asp231 in NM-R3), in the O_1_ state at 1 ms after light illumination (Figs. S10B and S11B) (31). Asn112 is 3.1 Å apart from the PSB in the resting state of KR2, which resembles the positional relationship of Asn98 and the PSB in the anion-depleted structure or the O_1_ structure model of NM-R3 (Fig. S10). This structural similarity might be a common feature of the ion pumping rhodopsins in the absence of an ion around the retinal. A superimposition of the resting state and the O_1_ intermediate of KR2 indicated that the Cα atom positions of Asn112 and Asp251 did not change very much (Fig. S11B). Therefore, the conformational changes in NM-R3 from the resting state to the O_1_ state differ from those of KR2 (Fig. S11).

## Conclusion

Our TR-SFX data clearly visualized dramatic conformational changes in NM-R3 after halide ion release (Fig. 4). The 13-*cis*/15-*syn* configuration of the retinal is displaced upward toward helix C, breaking the connectivity to the extracellular side. The upward displacement of the retinal induces the sequential movements of Trp99 and Trp201 toward the cytoplasmic side, to change the ion channel around the retinal. Asn98 is substantially shifted to stabilize the retinal binding pocket in the absence of the halide ion, accompanied with the large movements of helix C. Meanwhile, Asp231 of helix G approaches the Asn98 side chain, which also blocks the accessibility from the extracellular side. These structural changes in NM-R3 are similar to those of *Np*HR observed in low temperature trapping studies, but differ from those of KR2 or bR obtained from TR-SFX data. Therefore, the considerable structural changes may be specific to the rhodopsin chloride pump, which transports an anion with a relatively large ionic radius. In addition, the difference electron density map obtained from NM-R3 crystals with iodide ions suggests that the halide ion slightly moves toward Thr102 and the displacement of Asn98 to the halide binding site has already begun at Δ*t* =10 μs. Our observations of anomalous signals from iodide ions provide clues toward solving the pathway of halide ion transfer in NM-R3.

## Materials and Methods

### Detailed materials and methods are provided in *SI Appendix*

NM-R3 for crystallization was synthesized by using an *E. coli* cell-free protein synthesis system, and the protein was crystallized by the *in meso* method (20). The microcrystals were soaked in crystallization buffer with 1,000 mM NaBr.

TR-SFX experiments were performed at BL3 of SACLA (47). NM-R3 crystals were loaded into the injector (34). A 540 nm wavelength pump laser was introduced through optical fibers. Data collection was monitored by Cheetah and data processing was performed using CrystFEL (48, 49). The structure was solved by molecular replacement with PDB entry 5B2N (20), and difference Fourier maps were calculated by the CCP4 and PHENIX suites (50, 51).

The photocycle reaction of NM-R3 crystals with 1,500 mM NaBr crystallization buffer was induced by a 532 nm pump pulse, and the spectral changes during the reaction were measured with a spectrometer. The pump and probe light source and the spectrometer were synchronized by pulse generators, and the pump-probe delay time was adjusted with a timing jitter of ± 20 ns.

## Abbreviations

PDB: Protein Data Bank
DDM: n-dodecyl-b-_D_-maltopyranoside
rmsd: root-mean-square-deviation.

## Acknowledgements

We acknowledge members of the Engineering Team of RIKEN SPring-8 Center for technical support. We are grateful for computational support from the SACLA High Performance Computing System and the Mini-K supercomputer system. XFEL experiments were conducted at BL3 of SACLA, with the approval of the Japan Synchrotron Radiation Research Institute (JASRI) (proposal numbers 2017A8019, 2017A8028, 2017B8022, 2017B8023, 2018A8012, 2018A8023, 2018B8073, 2019A8013). This work was supported by MEXT/JSPS KAKENHI Grant Numbers 17K07324 and 20H05450 (T.Ho.), 19H05784 (M.K.), JP19H05776 (S.I.), and 18H02394 and 21H02439 (E.N.); and the Platform Project for Supporting Drug Discovery and Life Science Research (Basis for Supporting Innovative Drug Discovery and Life Science Research (BINDS)) from AMED under Grant Number JP21am0101070 (S.I.). We thank H. Fukuzawa for discussions on emissions from samples containing halide ions.

