## Supplementary Material for "Unidirectional ion transport mechanism of a light-driven chloride pump revealed using X-ray free electron lasers"

**This file includes**

| | |
|---|---|
| Supplementary Materials and Methods | pp. 2–6 |
| Supplementary Figures 1–12 | pp. 7–18 |
| Supplementary Tables 1–3 | pp. 19–21 |
| Supplementary References | pp. 22 |

### Supplementary Materials and Methods

#### Cell-free protein synthesis, purification, and crystallization of NM-R3

Cell-free protein synthesis of NM-R3 was performed essentially according to the previously reported protocols, with 10 mM magnesium acetate (1). After solubilization in 1.0% n-dodecyl-β-D-maltopyranoside (DDM), the supernatant containing NM-R3 was affinity purified on Ni-NTA resin with buffer A (20 mM HEPES (pH 7.4), 1 M NaBr, 0.04% DDM, and 400 mM imidazole). The affinity-tag was cleaved with TEV protease. The cleaved tag and TEV protease were removed by passage through the Ni-NTA column, and the NM-R3 protein was recovered from the flowthrough fraction. The protein solution was applied to a Superdex 200 16/600 column (GE Healthcare), equilibrated in 20 mM HEPES (pH 7.4), 1 M NaBr, 0.04% DDM and 1 mM dithiothreitol (DTT). The resulting peak fraction from the size-exclusion chromatography (SEC) was collected and concentrated with a 50 K-MWCO Amicon Ultra filter unit to 80 mg/ml.

A 12 µl portion of the NM-R3 solution were mixed with 18 µl of monoolein, using Hamilton syringes and a syringe coupler. After reconstitution into the Lipidic Cubic Phase (LCP), the syringe and coupler were removed and a clean wire (approximately 2 cm long) was inserted into the protein-laden LCP, according to the previously reported method for LCP-SFX (2). The protein-laden LCP (30 µl) was extruded as a straight column from a syringe and inserted into a 0.5 ml tube filled with precipitant solution (Fig. S12A). Crystals in the space group *C2* were grown in the dark at 20 °C in 38% PEG400, 300 mM K phosphate, and 100 mM HEPES (pH 7.0) (Fig. S12B).

For the preparation of crystals in the space group $P2_12_12_1$, the cell-free protein synthesis was the same as that of the *C2* crystal, except for the concentration of magnesium acetate (15 mM). In addition, the sample was finally purified by SEC on a Superdex 200 10/300 column, in 20 mM HEPES (pH 7.4), 1 M NaCl, 0.04% DDM and 1 mM (DTT). Crystals in the space group $P2_12_12_1$ were grown in 100 mM HEPES (pH 7.0), 400-700 mM K phosphate, and 24-30% PEG400 (Fig. S12C).

#### Sample preparation for LCP-SFX

The NM-R3 microcrystals stuck to the wire were soaked in 30% PEG400, 300 mM K phosphate, 100 mM HEPES (pH 7.0), 10 mM DTT, and 200 mM NaI or 1,000 mM NaBr for 12-24 hrs. The crystals were collected by grasping part of the wire with forceps prior to loading into a Hamilton syringe. After homogenizing the sample, a further 5 μl of liquid paraffin was added to 60 μl of LCP NM-R3 microcrystals and mixed for smooth flow.

**Experimental setup and X-ray free electron laser data collection**

TR-SFX experiments were performed at BL3 of SACLA (3, 4) in April 2017, July 2018, February 2019, and June 2019. NM-R3 crystals loaded into a high viscous sample injector (5) were kept at 20 ˚C during the measurement. A vacuum cleaning device was placed directly below the nozzle of the injector, to form a straight sample column (5). A 540 nm wavelength pump laser was introduced through optical fibers, as described below (6). The injector and the pump laser setup were mounted on motorized x-y-z stages for the intersection of the sample stream with the XFEL beam and the pump laser beam.

Data were collected using 7.0 keV X-ray pulses with a pulse duration of < 10 fs and a repetition rate of 30 Hz. NM-R3 crystals in the LCP matrix were extruded through a 75 μm diameter nozzle at a flow rate of 2.5 μl/min, an order of magnitude faster than the usual rate, to completely replace the sample region exposed to the visible laser pulse before the arrival of the next XFEL pulse. In total, 31,017 images for NM-R3-Br and 85,087 images for NM-R3-I were collected as a "complete dark" data set without the use of the pump laser during data collection.

**Pump laser setup for TR-SFX**

A nanosecond visible light pump pulse with a wavelength of 540 nm was supplied from the output of an optical parametric oscillator (NT230, Ekspla), which was controlled at 10 Hz with a pulse generator (DG645, Stanford Research Systems). The pulse generator was also used to synchronize the pump pulse with the XFEL pulse and control the delay time (Δ*t*), with a timing jitter of ±0.5 ns. A 50:50 beamsplitter was used to divide the 540 nm pump beam into two, with each half coupled into an optical fiber (100 μm core diameter, 0.22 numerical aperture) and delivered to the experimental setup. After collimation, the two pump beams were focused on the sample using objective lenses (33.5 mm working distance, 0.28 numerical aperture) in an almost counter-propagating geometry. Samples were thus irradiated from both sides of the LCP microjet, a geometry chosen to optimize the crystal excitation efficiency. The pump focal size was set to 40 μm (FWHM) and the pump energy was 6 μJ (3 μJ from each direction) at the sample point. A portion of the pump beam (5%) was isolated with a beam sampler, and its signal detected by a photodiode was used to monitor the laser status and to tag a diffraction image with "light-on". Since the pump repetition rate was one-third (540 nm) of that of XFEL, the "light-on" and the two "dark" datasets were collected sequentially.

**Data processing and structural analysis**

Datasets were collected over four beam times. Data collection was monitored by a real-time data processing pipeline (7) developed on Cheetah (8). Images with more than 20 strong spots were written as hits and processed by CrystFEL, version 0.6.1 or 0.6.3 (9). Spot finding for indexing by DirAx (10) was performed within CrystFEL by Zaefferer's algorithm (11) or the peakfinder8 algorithm (5). Spot finding algorithms and parameters were optimized for each beam time. The detector distance and spot finding parameters were optimized by a brute-force search. The detector metrology was refined by the geoptimiser program (12). The radii of integration and background masks were 3 and 7, respectively. The lattice constants of NM-R3 crystals exhibited indexing ambiguity characterized by an operator (h,-k,-h-l). This ambiguity was resolved by the ambigator program. Integrated intensities were merged by Monte Carlo integration with frame-wise scaling and resolution cutoff in process_hkl. Data collection statistics are summarized in Supplementary Table S2.

Difference Fourier maps were calculated with the CCP4 suite (13) and the PHENIX suite (14). Because ambigator only resolved the indexing ambiguity within each time point, but not between time points, merged intensities were reindexed by the REINDEX program in the CCP4 suite or the Reflection file editor in the PHENIX program when necessary. This is the same procedure as described previously (2). Structure factor amplitudes were then obtained by CTRUNCATE. Finally, difference

Fourier electron density maps ($|F_{obs}|^{light}-|F_{obs}|^{dark}$)·exp[i$\Phi_{calc}$] were calculated by FFT (15), using phases calculated from the refined ground state structure.

**Structure determination and refinement**

The NM-R3 ground state structure was determined by molecular replacement using Molrep (16), with PDB entry 5B2N as the search model. The NM-R3 ground structure was built manually in Coot (17). After several rounds of rebuilding and refinement with REFMAC5 (18) and PHENIX (14), the final ground state structures containing bromine or iodine atoms were recovered with $R_{work}$ and $R_{free}$ of 17.6 % and 20.8 % or 17.2 % and 20.5 %, respectively (Table S2).

Intermediate state structures with bromide at $\Delta t$ = 1 ms were refined with PHENIX (14). Atom position coordinates (x, y, z) and group occupancies of the conformers of an intermediate were refined, but not their B-factors. Waters at the negative peak positions in the difference Fourier map were removed from the conformers of an intermediate of the protein. Refinement statistics and crystallographic occupancies of the intermediate conformation can be found in Table S2. Groups of atoms corresponding to the strongest peaks in the difference Fourier electron density maps for all time points were selected (Table S3). The anomalous electron density map for NM-R3-I at each time point was calculated by SHELXC (19) and ANODE (20), using unmerged intensity data.

**Time-resolved spectroscopy**

The photocycle reaction was induced by a 6-ns, 532-nm pump pulse from an optical parametric oscillator (OPO) (NT230, EKSPLA), and the spectral changes during the reaction were measured with a fiber-coupled spectrometer (Flame-S, Ocean Optics) with a microsecond white-light probe pulse from a Xe-flash lamp (L11316-11-11, Hamamatsu Photonics). A portion of the probe light (5%) was isolated and used to correct the pulse-to-pulse fluctuations of the probe light intensity. The pump and probe lights were focused on the sample point with beam diameters of 300 μm and 40 μm, respectively. The pump and probe light sources and the spectrometer were synchronized by pulse generators (DG645, Stanford Research Systems), and the pump-probe delay time was adjusted with a timing jitter of ± 20 ns. The pump and probe repetition rates were 0.5 Hz and 1 Hz, respectively. The microcrystals were suspended in 100 mM

HEPES (pH 7.0) containing 30% PEG400, 300 mM potassium phosphate, and 1.5 M NaBr, and packed with two quartz windows with a 70 μm-thick spacer. The pump energy was adjusted to be 20 μJ at the sample point. The global exponential fitting analysis was performed by Igor Pro (WaveMetrics). In the analysis, the TR difference spectra $\Delta A(\lambda, t)$ were fitted with a sum of exponentials with apparent time constants $\tau_i$ and amplitudes $A_i(\lambda)$, as follows: $\Delta A(\lambda, t) = -A_1(\lambda)\cdot\exp(-t/\tau_1) - A_2(\lambda)\cdot\exp(-t/\tau_2)$. In the present case, $A_1(\lambda)$ corresponds to a difference of $[\Delta A_O(\lambda) - \Delta A(\lambda, t = 10\ \mu s)]$, whereas $A_2(\lambda)$ corresponds to $[\Delta A(t = 200\ ms) - \Delta A_O(\lambda)]$. Here, $\Delta A_O(\lambda)$ represents the TR difference spectrum of O minus NM-R3.

**Supplementary Figures**

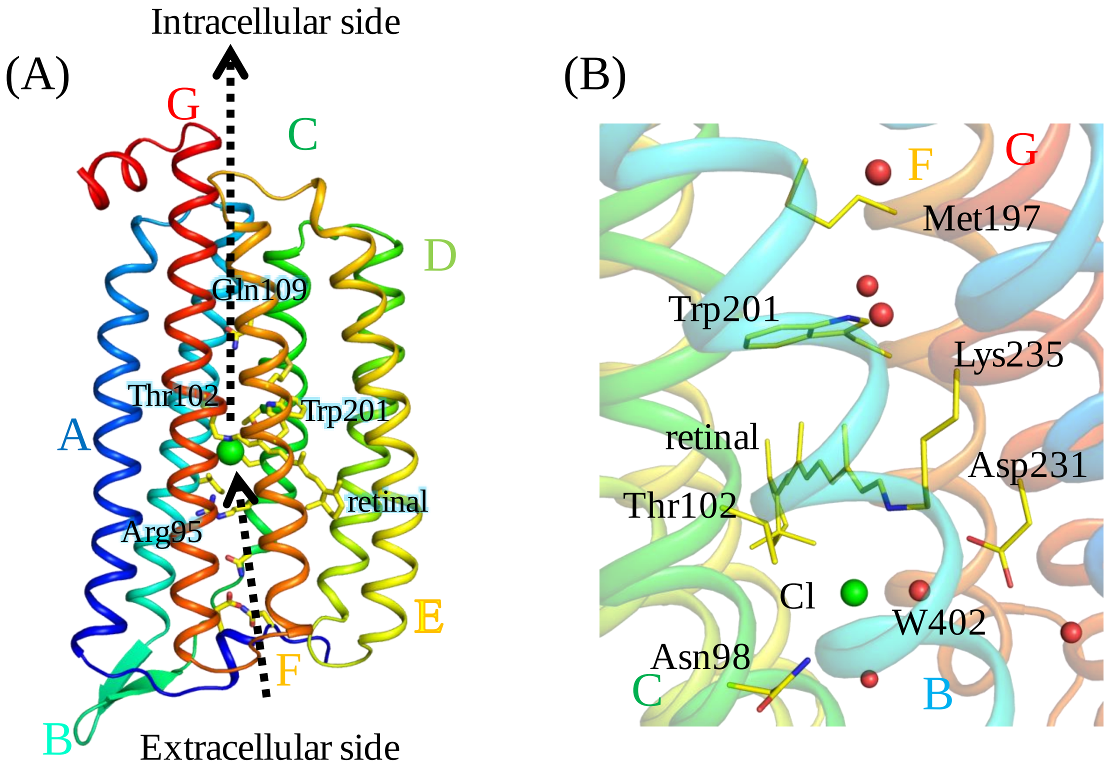


**Fig. S1. Structure of NM-R3.**
(A) Overall structures of NM-R3 with a chloride ion (PDB ID: 5B2N) (1). Transmembrane helices are labeled A to G. Black dashed arrows suggest the movement of chloride ions following photoexcitation. (B) Structures of the chloride binding region and the retinal of NM-R3. The chloride ion and water molecules are depicted by green and red spheres, respectively. The retinal and putative key residues for ion-transporting are shown in stick models.

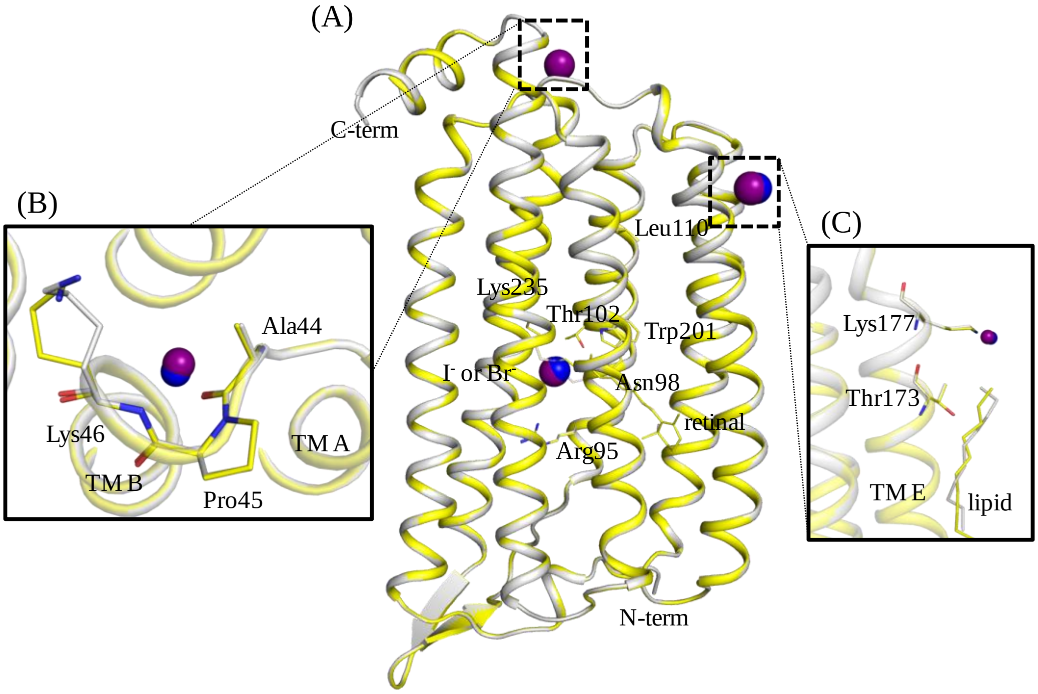


**Fig. S2. Comparison of the resting state structures of NM-R3-Br and NM-R3-I.** (A) Overall structures of the resting state NM-R3-I (gray) and the superimposed structure of NM-R3-Br (yellow). The stick models and black characters indicate the amino acid residues and retinal. Iodide and bromide ions are depicted as purple and blue spheres, respectively. (B) Second anion-binding site on the cytoplasmic loop of NM-R3. The anion (purple or blue spheres) is coordinated by Ala44, Pro45 and Lys46. (C) Third anion-binding site near TM3 of NM-R3. This anion (purple or blue spheres) is coordinated by Ser11 of the neighboring NM-R3 protomer in the crystal lattice.

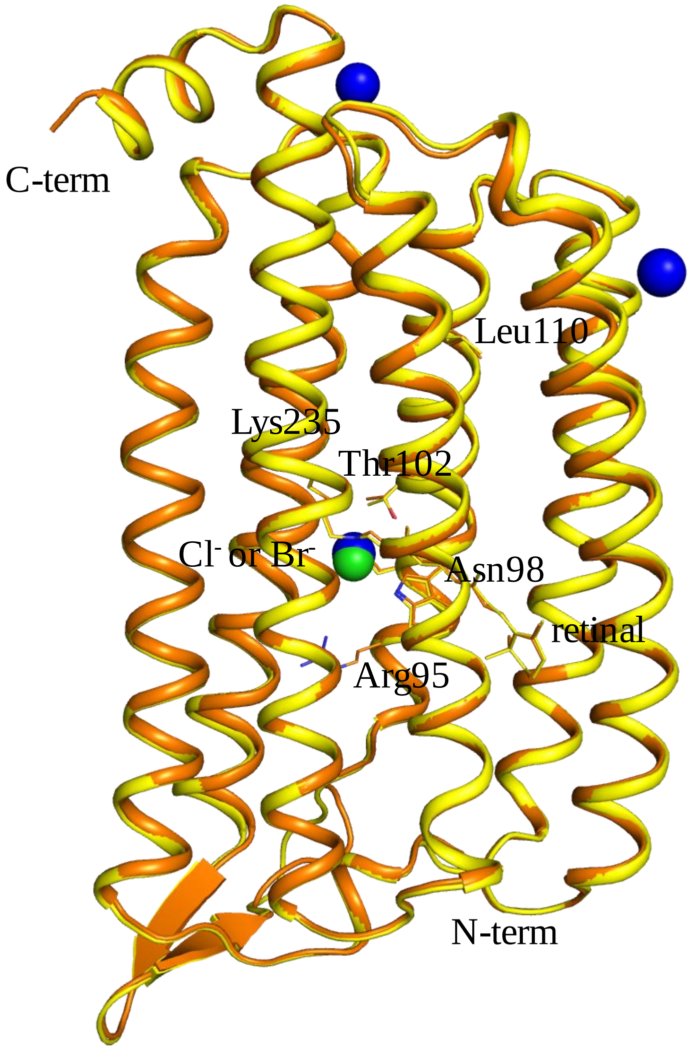


**Fig. S3. Comparison of the overall structures of NM-R3 from XFEL data and synchrotron data.**

Overall structures of the resting state NM-R3-Br (yellow) determined at XFEL and the superimposed structure of NM-R3-Cl determined from synchrotron data (orange, PDB ID : 5B2N) (1). The stick models and black characters indicate the amino acid residues and retinal. Bromide and chloride ions located near the retinal Schiff base are depicted as blue and light-green spheres in the structures, respectively.

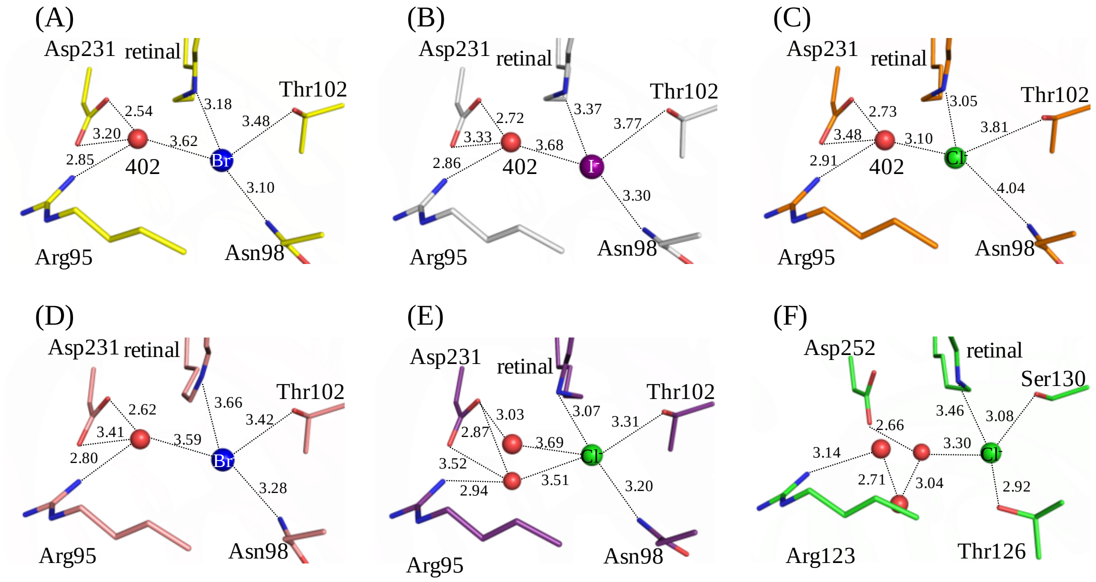


**Fig. S4. Comparison of the anion binding regions of various NM-R3 and *Np*HR structures.**

The anion binding site near retinal from the (A) SFX-structure of NM-R3-Br, (B) SFX-structure of NM-R3-I, (C) synchrotron-structure of NM-R3-Cl (PDB ID : 5B2N) (1), (D) synchrotron-structure of NM-R3-Br (PDB ID : 5G2A) (21), (E) SFX-structure of NM-R3-Cl (PDB ID : 5ZTL) (22), and (F) synchrotron-structure of *Np*HR (PDB ID : 3A7K) (23) are shown as stick models colored yellow, gray, orange, salmon, purple and green, respectively. Bromide, iodide, chloride, and water molecules located near the Schiff base of the retinal are depicted as blue, purple, green, and red spheres in the structures, respectively. Numbers and dashed lines indicate the distances (Å) between two atoms.

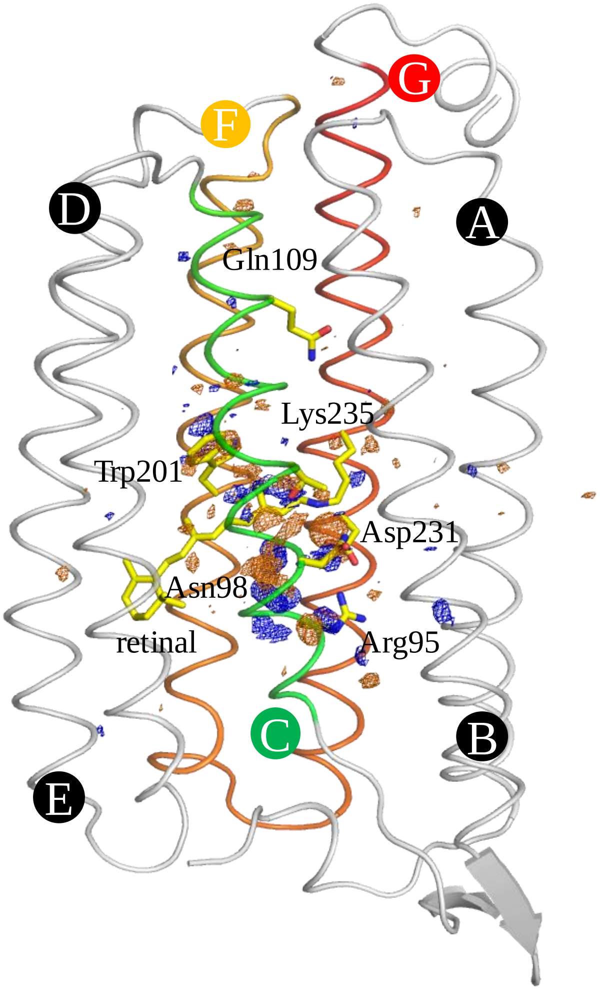


**Fig. S5 Overview of the $|F_{obs}|^{light}$ - $|F_{obs}|^{dark}$ difference Fourier electron density maps near the retinal of NM-R3-Br at Δ*t* = 1 ms.**
Blue represents positive difference electron density (contoured at +3.8σ) and orange is negative difference electron density (contoured at -3.8σ). The NM-R3-Br resting state structure was used for phases in calculating these difference Fourier maps and is shown in yellow. Transmembrane helices C, F, and G are colored green, orange, and red, respectively.

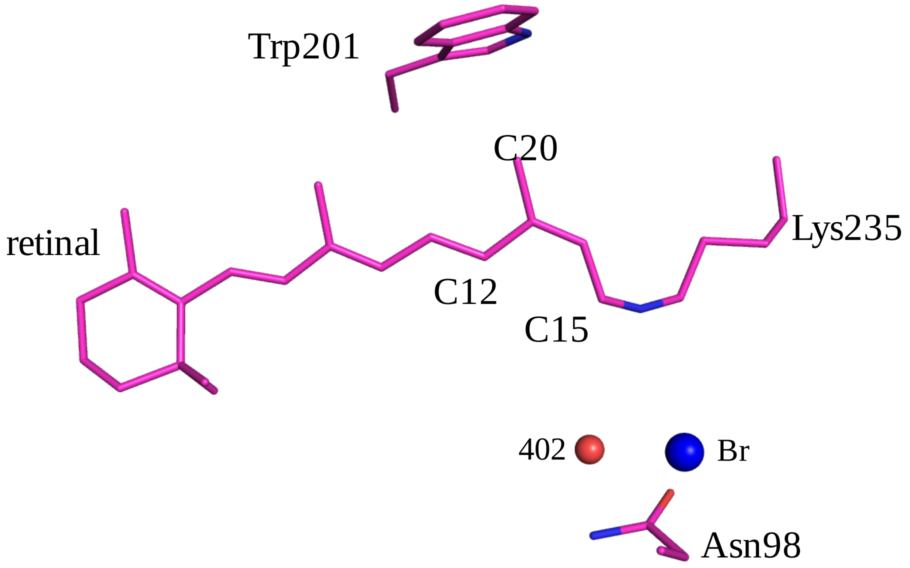


**Fig. S6. The anion binding site and the retinal of the $O_1$ intermediate structure at $\Delta t$ = 1 ms.**
Crystallographic structural models of the retinal and surrounding amino acids. The retinal adopts the 13-*cis*/15-*syn* form at $\Delta t$ = 1 ms (pink). The bromide ion and water molecule, Wat402, are depicted by blue and red spheres, respectively.

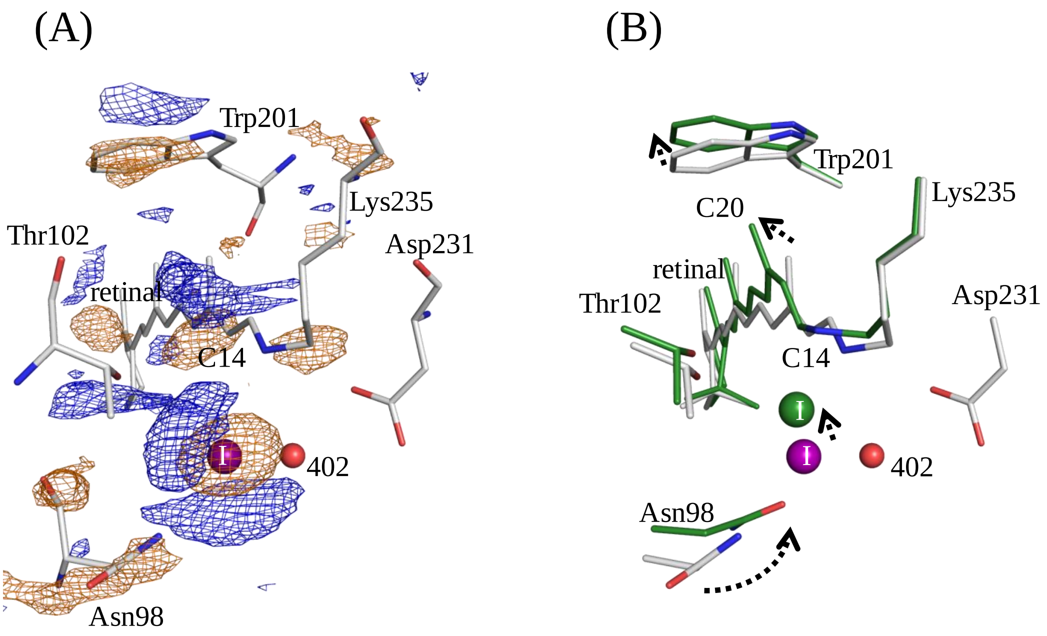


**Fig. S7. Difference map near the retinal in NM-R3-I at Δ*t* =10 μs.**
(A) View of the $|F_{obs}|^{light}$ - $|F_{obs}|^{dark}$ difference Fourier electron density map near the retinal for Δ*t* =10 μs. The blue electron density map indicates positive electron density, and orange denotes negative electron density (contoured at ±3.0σ). The resting state NM-R3 model (gray) was used for phases when calculating this map. (B) Crystallographic structural models derived from partial occupancy refinement are superimposed upon the resting state NM-R3 structure (gray) for Δ*t* = 10 μs (green). The movements of the molecules are depicted by dashed arrows. The iodide ion (in the resting state and Δ*t* =10 μs) and Wat402 are depicted by purple, green, and red spheres, respectively.

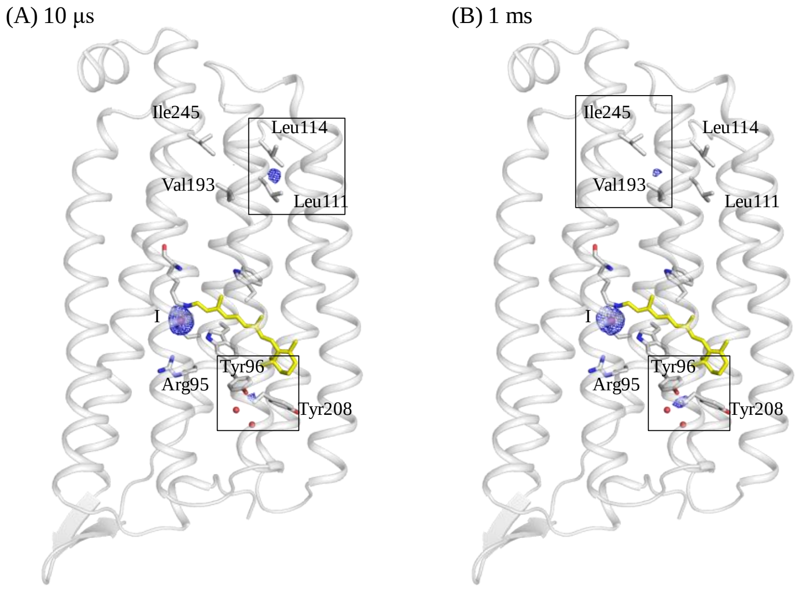


**Fig. S8. Anomalous density map at Δ*t* =10 μs and 1 ms.**
Anomalous density map (blue mesh) at Δ*t* =10 μs (A) or Δ*t* =1 ms (B), superimposed with the resting state structure model (gray). The anomalous peak region of NM-R3 is shown as a gray stick model. The map shows only anomalous peaks around the anion binding site and in the box. The map is contoured at 3.0σ.

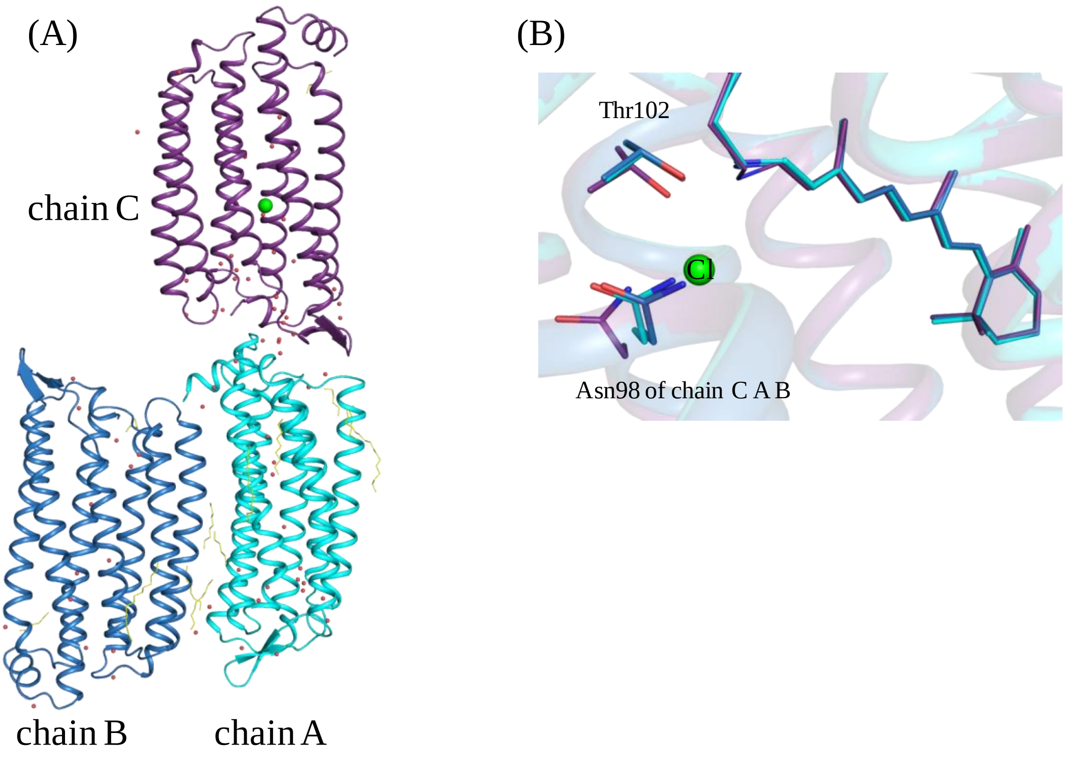


**Fig. S9. Overall structure of NM-R3 in the space group $P2_12_12_1$.**
(A) The crystal structure of NM-R3 at 2.3 Å resolution in the space group $P2_12_12_1$. The asymmetric unit contains three molecules, A, B, and C. (B) Chains A (cyan) and B (blue) are in the anion-depleted form, and chain C (purple) is in the chloride-bound form. Chloride ions and water molecules are depicted by light-green and red spheres, respectively.

(A) (B)

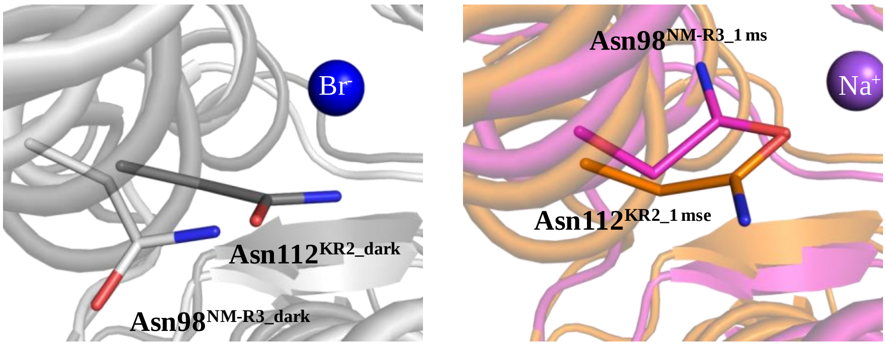


**Fig. S10. Conformational changes around Asn98 in NM-R3 and Asn112 in KR2.** (A) Superimposed structures of the resting state NM-R3 structure (gray) with the resting KR2 structure (dark gray, PDB ID = 6TK7) (24). (B) The superimposed structures of the $O_1$ intermediate NM-R3 (pink) structure with the $O_1$ intermediate KR2 structure (orange, PDB ID = 6TK2) at 1 ms after photoactivation (24). Bromide and sodium ions are depicted by blue and purple spheres, respectively.

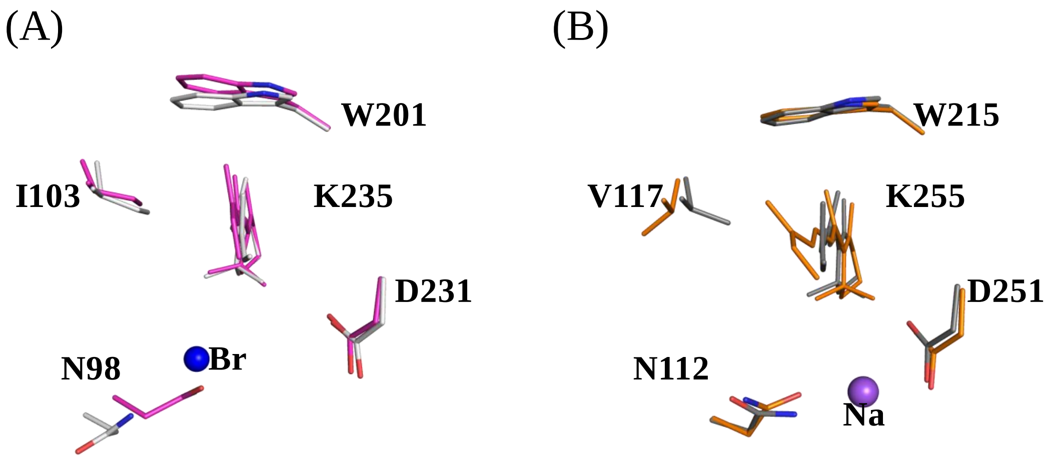


**Fig. S11. Conformational changes of the retinal binding sites in NM-R3 and KR2 at Δ*t* =1 ms.**
(A) Superimposed structures of NM-R3 for Δ*t* =1 ms (magenta) on the resting state structure (gray). (B) Superimposed structures of KR2 for Δ*t* =1 ms (PDB ID = 6TK2, orange) on the resting state structure (PDB ID = 6TK7, dark gray) (24). Bromide and sodium ions are depicted by blue and purple spheres, respectively.

(A) (B) (C)

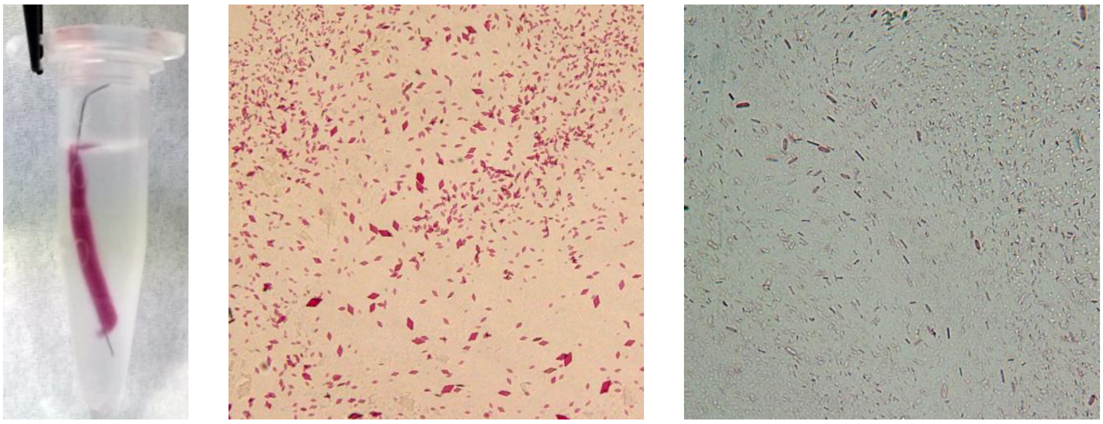

**Fig. S12**. LCP crystallization of NM-R3 using a thin wire.
(A) Protein-LCP with a wire in 0.5 ml tube. (B) Crystals in space group *C2*. (C) Crystals in space group $P2_12_12_1$.

**Supplementary Tables**

**Table S1. Partial alignments of functionally important amino acids of helices C, F, and G of each rhodopsin**

| | Function | helix C | | | | | helix F | | helix G | |
|---|---|---|---|---|---|---|---|---|---|---|
| NM-R3 | $Cl^-$ | R95 | N98 | W99 | T102 | Q109 | M197 | W201 | D231 | K235 |
| NpHR | $Cl^-$ | R123 | T126 | W127 | S130 | A137 | T218 | W222 | D252 | K256 |
| bR | $H^+$ | R82 | D85 | W86 | T89 | D96 | T178 | W182 | D212 | K216 |
| KR2 | $Na^+$ | R109 | N112 | W113 | D116 | Q123 | F211 | W215 | D251 | K255 |

The positions corresponding to N98, T102, and Q109 in NM-R3 are colored red.

**Table S2: Crystallographic data and refinement statistics.**

| | Br | | $P2_12_12_1$ |
|---|---|---|---|
| | Dark | 1 ms | Dark |
| PDB ID | 7VGT | 7VGU | 7VGV |
| ***Data collection*** | | | |
| Resolution rage | 42.8-2.1 (2.14-2.10) | | 46.3-2.3 (2.34-2.30) |
| Space group | *C2* | | $P2_12_12_1$ |
| Unit cell (Å) | a=104.3, b=51.1, c=78.4 | | a=68.4, b=69.5, c=231.4 |
| Unit cell (˚) | α=90, β=131.5, γ=90 | | α=90, β=90, γ=90 |
| Number of collected images | 256194 | 1120589 | 394606 |
| Number of hits | 35639 | 54265 | 74395 |
| Number of indexed patterns | 31017 | 46447 | 25936 |
| Indexing rate (%) | 87.0 | 85.6 | 34.9 |
| Number of total reflections | 18261 | 18261 | 50102 |
| Number of unique reflections | 18261 | 18261 | 50102 |
| Completeness (%) | 100 (100) | 100 (100) | 100 (100) |
| Multiplicity | 213 (56.3) | 363 (91.0) | 136 (95.0) |
| $R_{split}$(%) | 10.4 (32.5) | 8.24 (34.8) | 11.9 (95.6) |
| $CC_{1/2}$ | 98.8 (89.5) | 99.3 (91.8) | 98.4 (47.3) |
| I/σ(I) | 9.35 (4.0) | 11.6 (4.1) | 5.41 (1.2) |
| ***Refinement*** | | | |
| Number of refined atoms | 2206 | 242 | 6429 |
| Intermediate occupancy (%) | - | 19 | - |
| $R_{work}$ / $R_{free}$ (%) | 17.6/20.8 | 18.2/21.0 | 21.5/24.7 |
| Average B factor (Å$^2$) | | | |
| Protein | 32.1 | 31.5 | 53.2 |
| Retinal | 22.7 | 22.7 | 41.6 |
| Water molecules | 41.6 | 41.4 | 56.7 |
| Lipids | 53.6 | 52.1 | 60.2 |
| Ramachandran plot | | | |
| Favored | 99.2 | 99.2 | 99.2 |
| Allowed | 0.8 | 0.8 | 0.8 |
| Disallowed | 0 | 0 | 0 |
| r.m.s.d. bond lengths (Å) | 0.002 | 0.004 | 0.003 |
| r.m.s.d. bond angles (˚) | 0.499 | 0.640 | 0.536 |

**Table S3. Significant peaks in difference maps**

| | Bromide | Iodide | |
|---|---|---|---|
| | 1 ms | 1 ms | 10 μs |
| Anion | -18.77 | -22.01 | -18.10 |
| C11 - Lys Cε | +7.37 | +6.05 | +4.89 |
| C12 - C14 | -6.30 | -6.06 | -5.11 |
| NZ | NA | -3.29 | -4.83 |
| C20 | -5.58 | -4.97 | -3.34 |
| Arg95 | +5.46 | +4.25 | NA |
| Val97 | +6.66 / -6.75 | +5.25 / -6.09 | -4.21 |
| Asn98 | +7.01 / -5.57 | +5.98 / -3.80 | +9.28 / -4.46 |
| Trp99 | +5.79 / -5.41 | +5.25 / -6.04 | +4.09 / -4.70 |
| Met100 | +4.18 / -4.16 | +4.39 / -4.28 | -3.59 |
| Met197 | +4.24 / -4.08 | +3.74 / -4.18 | |
| Trp201 | +5.70 / -6.34 | +6.02 / -5.06 | +4.41 / -4.37 |
| Asp231 | +5.53 | +4.63 | +4.29 |
| Ser234 | +4.05 / -4.82 | +5.10 / -3.66 | -3.88 |
| Wat401 | -4.54 | -3.23 | NA |
| Wat402 | -4.15 | -3.20 | NA |
| Wat491 | +3.37 | NA | NA |
| Wat492 | +3.78 | NA | NA |
| Wat493 | +3.62 | NA | NA |
| Wat501 | -3.86 | -3.22 | NA |
